# Scalable high-density tactile arenas for measuring ground-contact forces in freely moving animals

**DOI:** 10.64898/2026.10.22.760735

**Authors:** Weiwen Lu, Ariel Slepyan, Pierce Perkins, Alessandro Ascani Orsini, Kiara Quinn, Nitish Thakor

## Abstract

Understanding how animals physically interact with their environment is fundamental to studying movement, sensorimotor control, and neurological function. Yet most behavioral measurement systems primarily capture movement and posture, while direct measurements of contact forces often require specialized instrumentation or involve trade-offs among spatial coverage, resolution, and acquisition speed. Here, we introduce a scalable tactile sensing platform that transforms the surface beneath freely moving animals into a high-resolution contact-imaging instrument. By combining modular sensor arrays with adaptive compressive tactile sampling, the platform provides 16,384 tactile sensing elements at 2-mm pitch across a 512 × 128-mm runway, with nominal 1 kHz acquisition. Rather than exhaustively scanning the entire array, the system concentrates measurements around active contacts, enabling high-speed sensing across an extended behavioral surface. Synchronized overhead video and a camera-assisted analysis pipeline enable paw-resolved measurements of contact loading, stance dynamics, footprint morphology, interlimb weight redistribution, and center-of-pressure trajectories. In a longitudinal rat model of sciatic nerve injury, the platform detected reduced loading of the injured hindlimb and increased loading of the contralateral limb, including changes that conventional gait timing measures underestimated. The modular architecture, together with openly available hardware designs, software and data, provides an accessible frame-work for studying the mechanical interactions underlying freely moving behavior, with potential applications across neuroscience, injury, and motor recovery research.

## 1 Main

Understanding how animals interact with their physical environment is fundamental to the study of behavior, motor control, and neurological function. Animal movement is commonly studied through video, motion tracking, and other kinematic measurements, which reveal where an animal moves, how its body is positioned, and when its limbs contact the ground. However, observing movement alone does not fully describe the mechanical interactions between an animal and its environment. Two animals may exhibit similar movements while distributing their weight very differently, and an animal may continue to place an injured limb on the ground while substantially reducing the load it bears [1]. Direct measurements of ground contact and loading therefore provide an important complementary dimension of behavior, with particular relevance to neuroscience, musculoskeletal research, pain, aging, rehabilitation, and recovery from injury [2–4].

Existing approaches to rodent locomotor analysis include optical footprint systems, instrumented walkways, treadmills, and video-based tracking methods [5–11]. These technologies have substantially advanced quantitative behavioral analysis, providing measurements of stride length, stance and swing duration, paw placement, coordination, and limb kinematics [12]. Optical systems such as CatWalk and MouseWalker capture footprint geometry, timing and coordination in detail [13–15], and GAITOR provides an open-source framework for gait analysis [16]. Optical imaging has also shown that paw contact patterns change during stance [14], but these systems infer contact from the brightness of light scattered by frustrated total internal reflection (FTIR), which indicates paw loading only indirectly rather than as a calibrated force measurement [17]. Pressure-sensitive walkways and force plates, in contrast, quantify weight-bearing and ground reaction forces directly [4, 8, 18]: they have characterized limb-force coordination [19], detected compensatory loading after neurological injury [1] and enabled inexpensive, open-source monitoring of motor activity [20]. However, force plates provide little spatial information within individual contacts, and although commercial pressure walkways now reach millimeter-scale resolution in rodents [21], dense sensing architectures face trade-offs among sensing area, spatial resolution, acquisition speed, and hardware complexity [6, 7, 22, 23]. These limitations make it difficult to capture how mechanical interactions evolve across multiple contacts and throughout freely moving behavior.

A particularly valuable measurement is the distribution of normal ground-contact forces, which describes how an animal supports and transfers its weight during locomotion. Unlike paw placement or stance duration alone, contact loading can reveal asymmetries, compensatory weight redistribution, and changes in mechanical support that may accompany pain, weakness, neurological dysfunction, or recovery [1, 19, 24, 25]. Spatially resolved measurements provide additional information about how loading changes within an individual paw, including shifts in contact area and center of pressure (COP) throughout stance. Capturing these dynamics across an extended behavioral surface requires a sensing system that combines large-area coverage, high spatial density, and rapid acquisition. The challenge is therefore not simply to measure forces at isolated locations, but to make distributed contact-force measurement a practical and accessible part of behavioral observation.

Recent advances in high-density tactile sensing and compressive acquisition architectures offer a route toward this capability [26–29]. In conventional scanning architectures, increasing the number of sensing elements increases the measurements required to acquire a complete frame, creating a trade-off between sensing area, spatial resolution, and acquisition speed [23]. Adaptive tactile sampling can reduce this burden by exploiting the spatial sparsity of physical contact: during locomotion, most of the sensing surface remains unloaded, while meaningful pressure signals are concentrated beneath the paws. By directing measurements toward active contact regions while surveying the remaining surface, adaptive acquisition makes high-speed, high-density sensing practical across larger areas [29]. This creates an opportunity to extend tactile sensing from localized measurement devices to instrumented experimental environments, in which the surface itself records an animal’s physical interactions during movement.

Here, we introduce a scalable, high-density tactile sensing platform for measuring ground-contact loading in freely moving animals. As an initial demonstration, we construct a 512 *×* 128-mm runway comprising 16,384 tactile sensing elements at 2-mm pitch, with nominal 1-kHz acquisition. The platform combines modular piezoresistive sensing arrays with an adaptive sampling strategy that prioritizes active contacts while maintaining background surveillance of the surface. Rather than measuring every taxel in each frame, the system acquires a sparse subset of measurements and resolves detected contacts at full spatial density. This is the first rodent gait platform to combine millimeter-scale spatial resolution with adjustable kilohertz-rate acquisition across extended, highly customizable sensing surfaces. Its modular architecture also allows the sensing surface to be extended or reconfigured by combining additional sensor modules. Synchronized overhead video provides anatomical context for the tactile measurements, while a camera-assisted analysis pipeline identifies individual paw contacts and extracts loading, stance dynamics, contact morphology, interlimb imbalance, and COP trajectories. We demonstrate the method in freely walking rats and a longitudinal sciatic nerve pinch injury model [30, 31]. We characterize the sensing platform, examine the effects of temporal sampling on loading measurements, and evaluate camera-assisted paw assignment against manually annotated contacts. Finally, we show that after sciatic nerve pinch, direct contact measurements detect reduced loading of the injured hindlimb, a shift of load toward the contralateral limb, and changes in paw-contact morphology that are not fully captured by conventional gait timing measures. Together, these results establish a scalable approach to high-resolution contact imaging during freely moving behavior and illustrate its potential as a general-purpose measurement technology for experimental animal research. Hardware design files, firmware, analysis software, the recordings and a build guide are openly available (https://tactilearena.github.io).

## 2 Results

### 2.1 A scalable tactile sensing surface enables high-resolution contact imaging

We developed a modular tactile sensing platform that converts an extended walking surface into a spatially resolved contact-measurement instrument (Fig. 1A1,A2). The demonstrated system comprises four 64 *×* 64 piezoresistive sensing modules arranged as a 64 *×* 256 array, providing 16,384 tactile sensing elements at 2-mm pitch across a 512 *×* 128-mm runway. The array is protected by a waterproof walking surface, allowing animals to move freely over the sensors without tethering or body-mounted instrumentation. Four independent readout boards acquire tactile measurements, while synchronized overhead video provides complementary information about body position and orientation. Together, these measurements capture where an animal contacts the ground and how loading is distributed across those contacts during locomotion.

**Fig. 1.**
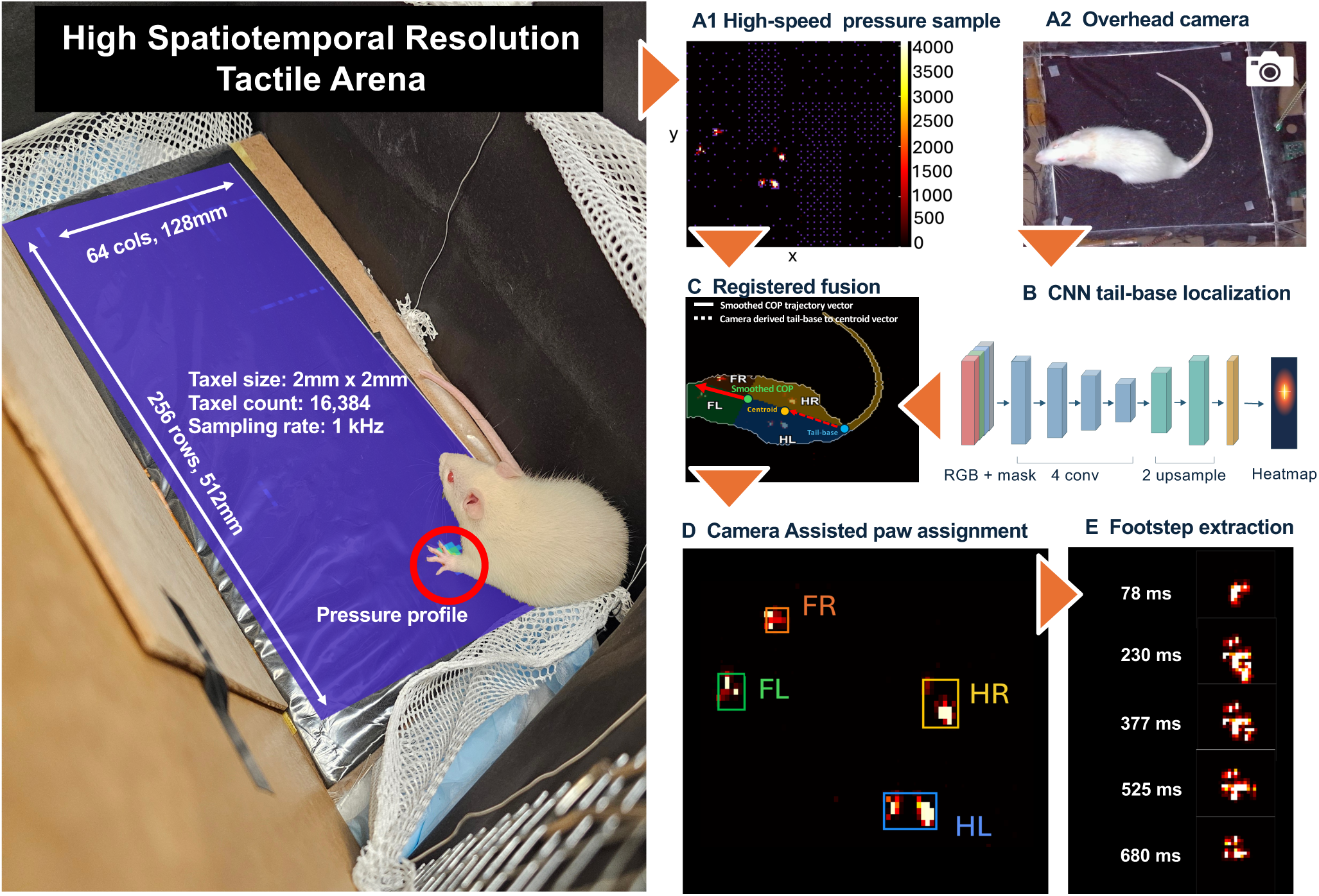
High-density tactile runway and camera-guided analysis. Four 64 *×* 64 sensor modules provide 16,384 tactile elements at 2-mm pitch. **A1,** Representative tactile frame acquired with adaptive quadtree sampling. Each module measures around 4.9% of its elements per frame: a coarse-to-fine background scan monitors the array, and any reading above threshold triggers full-resolution sampling of the surrounding neighbourhood, capturing each paw contact (bright regions). Black pixels indicate locations that were not measured in that frame, whereas purple pixels indicate sampled locations below the predefined contact threshold. **A2,** Overhead video. **B,** A convolutional encoder–decoder predicts a tail-base heatmap from an RGB frame and body mask. **C,** The body mask and predicted tail-base location are mapped into tactile coordinates and combined with the pressure data. Smoothed COP motion provides the walking-direction reference, while the tail-base–centroid vector supplies camera-derived body orientation. **D,** Contact positions relative to the smoothed COP, camera-derived paw votes, and temporal overlap constraints guide assignment to the left forepaw (FL), right forepaw (FR), left hindpaw (HL), and right hindpaw (HR). **E,** Successive pressure-contact detections are linked into paw-specific footstep events within manually curated walking windows.

The large sensing area, spatial density, and high acquisition rate are enabled by adaptive tactile sampling rather than exhaustive scanning of the full array [29]. Each 64*×*64 module contains 4,096 addressable taxels, but acquires a fixed budget of 200 measurements per frame, corresponding to approximately 4.9% of the module. Previously active regions are prioritized to maintain contact tracking, while a coarse-to-fine background scan surveys the remaining surface. When a measured taxel exceeds a contact threshold, neighboring taxels are sampled recursively to resolve the detected contact at full spatial density, subject to the measurement budget (Fig. 1A1; Supplementary Video 4). This approach concentrates acquisition on the relatively small regions where contact occurs, allowing nominal 1kHz acquisition across the runway without individually measuring all 16,384 taxels in every frame. Because each module operates with its own readout electronics, the architecture can be extended or rearranged to accommodate other sensing areas and experimental geometries.

The resulting sparse, indexed measurements are assembled into spatially registered tactile maps and synchronized with overhead video. A convolutional encoder–decoder identifies the animal’s tail base from the video image and body mask (Fig. 1B). The video-derived body mask and tail-base location are mapped into tactile coordinates, providing orientation cues alongside the pressure-derived center-of-pressure (COP) trajectory (Fig. 1C). Within manually curated walking windows, individual pressure contacts are detected and tracked through time. Contact locations relative to smoothed COP motion provide the primary basis for paw assignment, supplemented by camera-derived orientation information and temporal constraints. Contact fragments are then linked into paw-specific stance events (Fig. 1D,E). This combined measurement and analysis workflow connects large-area tactile imaging with anatomically interpretable contact dynamics during freely moving behavior.

### 2.2 Pressure sensor characterization

We assessed the tactile response during presses and controlled loading with an independent force plate (Supplementary Fig. S1). Across four consecutive presses, the spatially summed tactile signal followed the broad loading and unloading phases of the force-plate response. The root mean squared error (RMSE) between independently normalized signals was 0.101 over the active press intervals (Supplementary Fig. S1c). Because both signals were independently min–max normalized, this RMSE is unitless and amounts to about 10% of each signal’s full range; for scale, the representative press in Supplementary Fig. S1b peaked near 2 kg. Absolute calibration errors in grams are shown in Supplementary Fig. S1e. Pressure-map snapshots showed the formation, expansion, and disappearance of the contact region during a representative press (Supplementary Fig. S1b).

Board-specific calibration curves converted summed taxel responses to mass-equivalent loading (Supplementary Fig. S1d). Within each board’s recording, calibration was evaluated by withholding each selected press in turn and predicting its load from the remaining presses (Supplementary Fig. S1e).

### 2.3 Pressure imaging captures paw contacts during free locomotion

We next examined the pressure recordings obtained as rats moved freely along the runway. Projecting detected contacts onto synchronized video frames showed their positions relative to the animal throughout a representative walking sequence (Fig. 2a; Supplementary Videos 2 and 3). Individual pressure clusters resolved distinct contact patterns for each paw, including a representative left hindpaw contact recorded after surgery (Fig. 2b). Across a walking window, successive contacts formed a footprint trail that could be viewed alongside the geometric centroid derived from the overhead video (Fig. 2c). The corresponding paw-contact intervals captured the sequence and duration of contacts across all four limbs (Fig. 2d). Together, these examples illustrate the spatial and temporal information available from a single pressure-video recording.

**Fig. 2.**
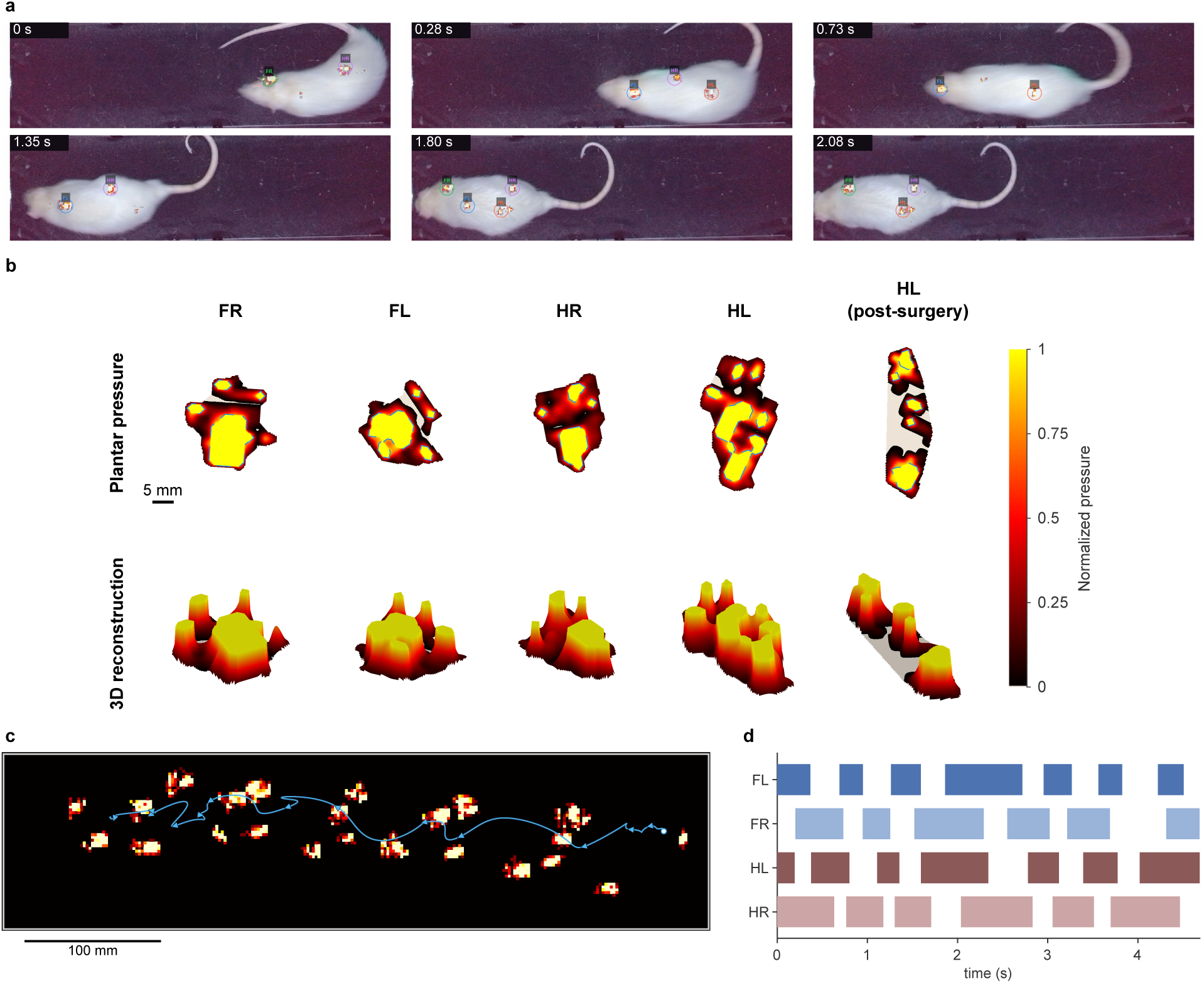
**a,** Detected paw contacts projected onto synchronized overhead video frames; labels indicate anatomical paw identity. **b,** Representative normalized plantar pressure maps and surface plots for each paw, including a representative postoperative HL contact whose elongated shape is consistent with increased heel contact. Color and surface height encode taxel values normalized to each contact’s own peak, so the panels show contact shape and do not compare loading magnitude; the postoperative HL contact is included to illustrate shape. The scale bar corresponds to 5 mm. **c,** A trail of detected footprints at their maximum contact area during a walking window, overlaid with the trajectory of the geometric centroid of the body mask extracted from the corresponding video recording. **d,** Contact intervals for all four paws during a representative walking sequence; bars indicate detected stance intervals.

### 2.4 COP tracking resolves locomotor trajectories and local loading shifts

The center of pressure (COP) describes how a rat distributes its weight across the supporting limbs, which the geometric centroid of the body in the camera image does not capture; this makes it particularly valuable for longitudinal studies of loading changes after injury. In a representative recording, the COP trajectory describes traversals of the runway (Fig. 3a). The corresponding longitudinal position and COP-speed traces describe movement along the runway and the temporal variation in COP motion within three selected walking windows (Fig. 3b,c).

**Fig. 3.**
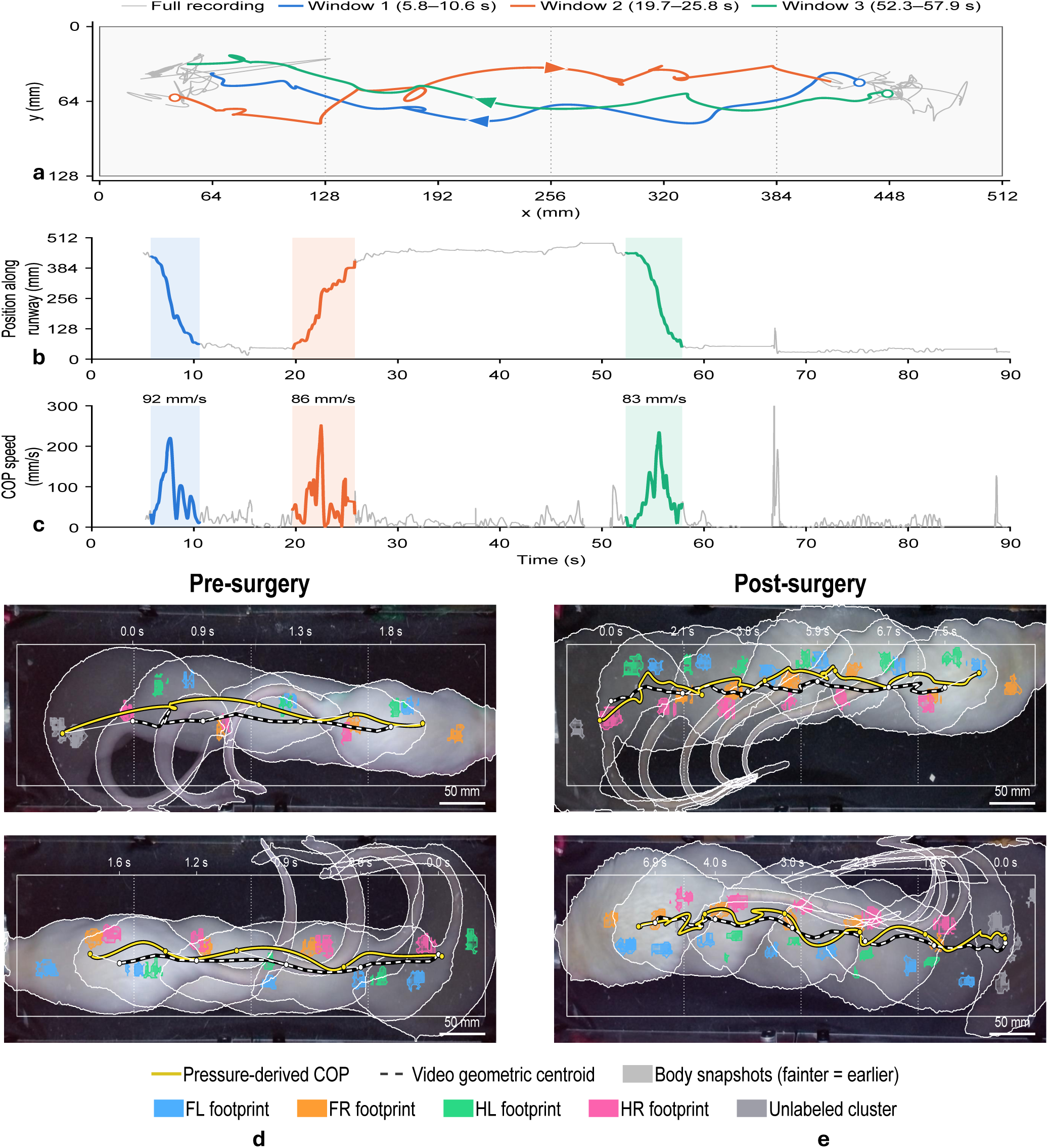
Whole-animal COP tracking and comparison with video-derived body position. **a,** Whole-animal COP trajectory over a representative recording (gray), with three selected time windows highlighted in blue (5.8–10.6 s), orange (19.7–25.8 s), and green (52.3–57.9 s). Arrows indicate travel direction. **b,** Longitudinal COP position over time, with the same windows highlighted. **c,** COP speed over time. Numerical annotations on the selected windows indicate the displayed average speed for the selected windows. COP speed describes motion of the pressure-weighted centroid, including redistribution of loading between contacts. **d,e,** Whole-animal COP (yellow solid lines) and video-derived body centroid (white dashed lines) in illustrative presurgical (**d**) and postsurgical (**e**) walking windows for the SNI group. The footprints are extracted from the peak of their corresponding footstep. The brightness of body mask indicates relative time within each window. Note that the windows differ in duration: the postsurgical windows in **e** last roughly four times longer than the presurgical windows in **d** for a similar distance traveled. The two trajectories don’t necessarily overlap, because the geometric centroid derived from the video is dependent on lighting condition as well as filter intensity, whereas the COP trajectory depends on the distribution of weight loading among the paws.

Whole-animal COP trajectories followed the movement of the video-derived body centroid in illustrative presurgical and postsurgical walking windows (Fig. 3d,e). In these selected examples, the postsurgical COP trajectories appeared more irregular, with more pronounced local curling and changes in direction than the presurgical trajectories. The postsurgical examples also took roughly four times longer to cover a similar distance and contained more paw contacts along the way, suggesting that the slower traversal reflects a reduced stride length rather than a lower cadence alone; this is consistent with the lower postsurgical COP speeds across walking windows (Fig. 4c). These observations suggest changes in weight-transfer patterns following injury, although further quantitative analysis is needed to determine whether they are consistent across animals. The platform is also capable of extracting COP trajectories for individual paw contacts, which provides more insights into changes in load shifting during stance for any particular foot(Supplementary Fig. S3).

**Fig. 4.**
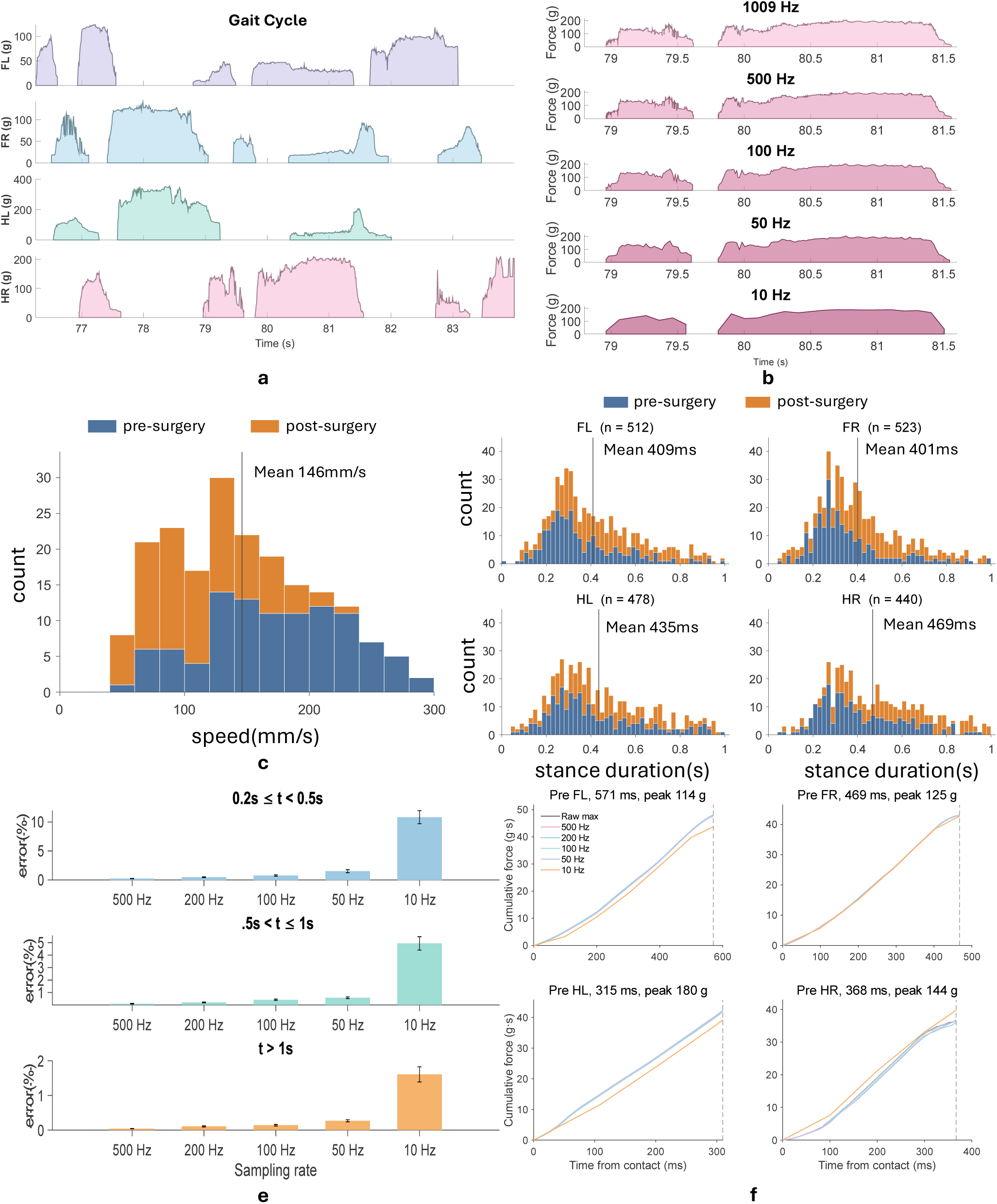
**a,** Paw-specific loading traces during a representative walking gait cycle. The plotted gram scale represents mass-equivalent loading rather than pressure units. **b,** A native 1,009-Hz loading trace and versions downsampled to 500, 100, 50, and 10 Hz. **c,** Distribution of COP speed across walking windows before and after surgery. COP speed describes pressure-weighted centroid motion rather than body walking speed. **d,** Distribution of stance duration for each paw across the same walking windows before and after surgery. **e,** Final cumulative-loading error relative to the native trace for contacts lasting 0.2–0.5 s, 0.5–1 s, and more than 1 s, as grouped in the plot. Note that the y-axis range differs between the three groups: the 10-Hz error falls from about 10% for the shortest contacts to under 2% for contacts longer than 1 s. **f,** Cumulative-loading curves for representative contacts from all four paws. Each panel has its own y-axis range. Counts in **c** refer to walking windows and counts in **d** to stance events, not independent rats.

### 2.5 Temporal downsampling increases cumulative-loading error for short contacts

We examined how temporal sampling affects paw-loading measurements by downsampling native recordings to lower effective rates. The representative trace was acquired at 1,009 Hz (Fig. 4b). COP-speed and stance-duration distributions describe pressure-centroid motion and contact timescales in the analyzed recordings (Fig. 4c,d). COP speed reflects motion of the pressure-weighted centroid, including shifts in loading between contacts. Stance duration determines the number of temporal samples available to describe each contact.

Paw-specific traces captured loading and unloading during stance (Fig. 4a). Downsampling reduced waveform detail, with the largest visible distortion at 10 Hz (Fig. 4b). In the summary grouped by durations, cumulative-loading error was greatest for the shortest contacts and increased as sampling rate decreased (Fig. 4e). Representative contacts likewise showed larger deviations in cumulative loading at 10 Hz than at the higher tested rates (Fig. 4f). These results support selecting acquisition rates according to contact duration and the measurement of interest. This analysis evaluated loading traces and their time integrals, not COP trajectories.

### 2.6 Camera-assisted decoding achieves high paw-assignment accuracy during walking

Paw identities were assigned from pressure-contact positions relative to smoothed whole-animal COP motion, with camera-derived paw votes providing supplementary orientation information. The tail-base localizer that supplies this camera-derived orientation reached a cross-validated median error of 6.6 mm (range across repeats 6.0–8.0 mm), with 72% and 90% of 232 labeled frames within 10 and 20 mm (Supplementary Fig. S2b,c). Within manually curated walking windows from 81 recordings, the pipeline correctly identified 2,303 of 2,469 manual-labeled contacts (93.3%; Supplementary Fig. S2d). Of these reference contacts, 2,406 were matched to an automatically labeled contact, giving 95.7% label agreement among matched contacts; 63 were unmatched and 103 were assigned an incorrect paw identity. This evaluation measures contact matching and paw assignment within manually curated walking windows, rather than automatic selection of walking windows.

### 2.7 Longitudinal measurements reveal altered hindlimb loading after sciatic nerve pinch

We evaluated longitudinal changes in three rats with sciatic nerve pinch injury (SNI) on the left hindlimb and two sham-operated rats. Recordings were collected before surgery and over the following two weeks. Sham recordings provided a comparison for surgical handling and repeated testing, although the small cohort limits separation of injury-specific effects from other sources of variation. Paw-loading measurements were normalized by body mass and expressed relative to each animal’s presurgical baseline (Fig. 5e,f).

**Fig. 5.**
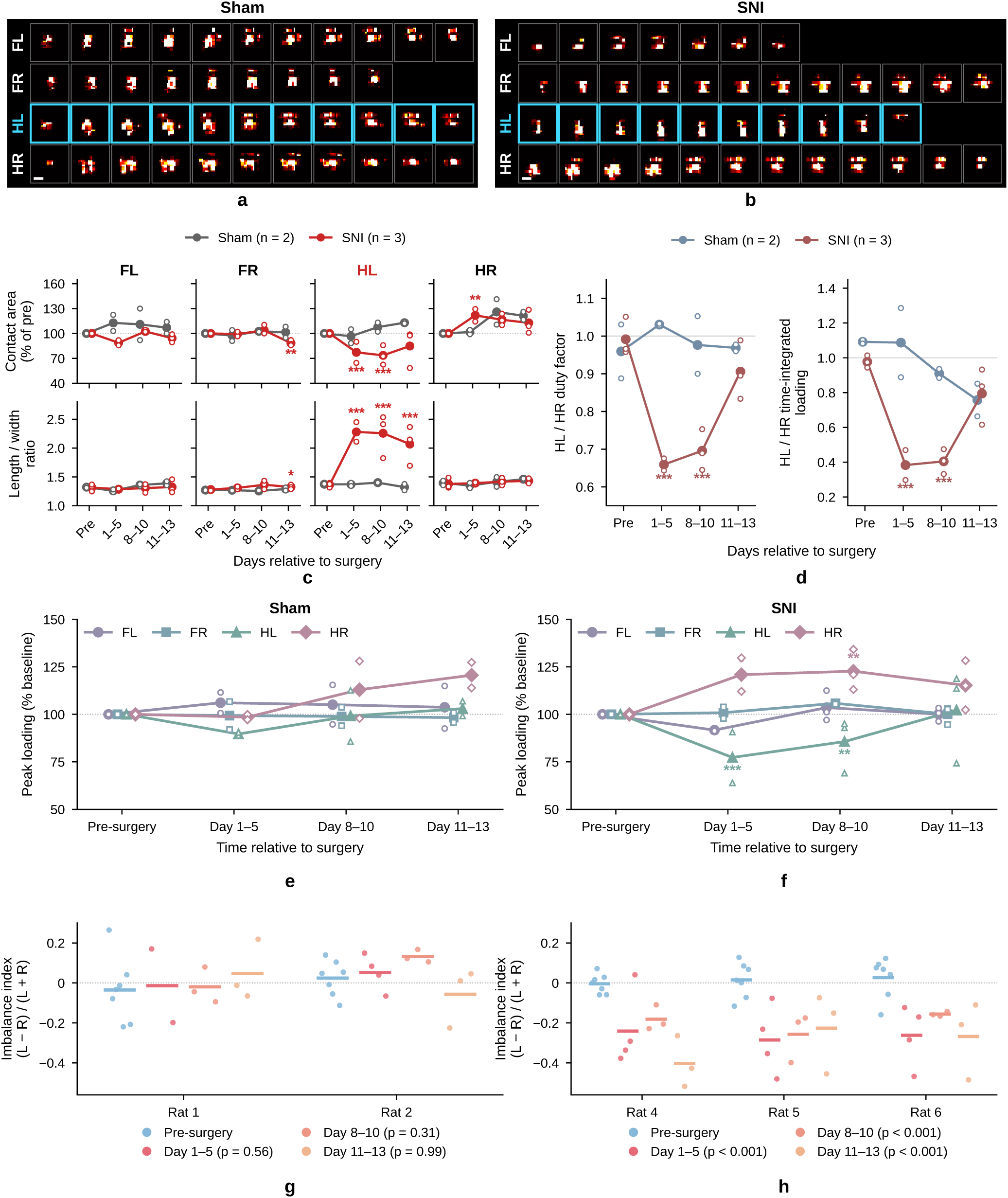
**a,b,** Successive pressure maps illustrate contact-shape evolution during a postoperative gait cycle in a sham-operated rat (**a**) and a rat with sciatic nerve pinch injury (SNI; **b**). Rows show the four paws; cyan outlines highlight the left hindpaw (HL). The illustrated injured HL contact appears more elongated than the sham HL contact. **c,** Contact area relative to baseline and footprint length-to-width ratio for each paw in sham (Rats 1–2) and SNI (Rats 4–6) rats. **d,** Left-to-right hindpaw duty-factor ratio (left) and time-integrated loading ratio (right). Walking-window ratios were median HL divided by median HR. **e,f,** Body-mass-normalized peak loading, expressed relative to each rat’s presurgical median for that paw, in sham (**e**) and SNI (**f**) rats. Colors and marker shapes identify paws. In **c–f**, small open markers show individual-rat values and lines with filled markers show group means; on days 1–5, two SNI rats contributed because Rat 4 remained mostly stationary. **g,h,** Hindpaw imbalance, (*L − R*)*/*(*L* + *R*), in sham (**g**) and SNI (**h**) rats. *L* and *R* are summed taxel responses within the labelled left and right hindpaw contacts during hindpaw-only support; positive values indicate greater left-sided loading. Points denote recording-session means and horizontal bars denote each rat’s epoch mean; all three SNI rats contributed at every epoch. Statistics (**c–h**): within each group, each postoperative epoch was compared with presurgical baseline using linear mixed-effects models fitted to recording-session means with a random intercept for rat and Satterthwaite degrees of freedom. Stars (**c–f**; colored by group in **c,d**) and legend *p*-values (**g,h**): *^∗^p <* 0.05, *^∗∗^p <* 0.01, *^∗∗∗^p <* 0.001.

Representative postoperative stance sequences illustrate the evolution of paw contacts in sham-operated and SNI rats, with a more elongated HL contact in the illustrated SNI example (Fig. 5a,b).

Footprint morphology changed most clearly in the injured left hindlimb (Fig. 5c). In SNI rats, HL contact area decreased to about 77% and 74% of baseline on days 1–5 and 8–10 (both *p <* 0.001) and was 85% of baseline on days 11–13 (*p* = 0.064), and the HL length-to-width ratio increased by 49%, 56% and 40% across the three postoperative epochs (all *p <* 0.001). The contralateral HR contact area increased to about 122% of baseline on days 1–5 (*p* = 0.009), and FR contacts were smaller and slightly more elongated on days 11–13 (area 88% of baseline, *p* = 0.003; length-to-width ratio +7%, *p* = 0.011). No morphology measure changed significantly in sham rats (all *p ≥* 0.05).

Hindpaw timing and loading ratios decreased during the early postoperative period in the SNI group (Fig. 5d). Relative to presurgical baseline, the duty-factor ratio decreased by 0.33 on days 1–5 and 0.30 on days 8–10; the corresponding decreases in the time-integrated loading ratio were 0.59 and 0.54 (all *p <* 0.001). By days 11–13, both ratios had moved toward baseline and no longer differed significantly from their presurgical values (duty factor, *p* = 0.305; time-integrated loading, *p* = 0.918). Neither ratio differed significantly from baseline in the sham group at any postoperative epoch (all *p ≥* 0.321).

Paw-specific peak-loading analysis localized the early loading changes to the hindpaws (Fig. 5e,f). In SNI rats, the loading of the injured left hindpaw decreased by 27% on days 1–5 (*p <* 0.001) and 18% on days 8–10 (*p* = 0.026), while the right hindpaw loading increased by 22% (*p* = 0.019) and 23% (*p* = 0.006), respectively. These percentages were obtained by back-transforming the log-scale model contrasts; the plotted curves summarize individual-rat medians relative to baseline. On days 11–13, neither hindpaw differed significantly from baseline (HL, *p* = 0.865; HR, *p* = 0.462). Forepaw loading in SNI rats and all paw loadings in sham rats showed no statistically significant postoperative differences (all *p ≥* 0.154). These comparisons support early unloading of the injured hindpaw with increased contralateral loading; the absence of a significant late difference does not establish equivalence to baseline. To examine how load was distributed between the hindlimbs in upright postures such as rearing, we analyzed frames in which only the hindpaws contacted the floor. For each session, we calculated a hindlimb imbalance index, (*L − R*)*/*(*L* + *R*), where negative values indicate right-sided loading. After surgery, the index shifted toward right-sided loading in all three SNI rats (Fig. 5h): relative to baseline, the index decreased by 0.28, 0.21 and 0.31 on days 1–5, 8–10 and 11–13 (all *p <* 0.001), so that the injured left hindpaw carried about 35–40% of the hindlimb load, compared with about 50% before surgery. Sham rats showed no significant change (all *p ≥* 0.31; Fig. 5g). Taken together, the data are consistent with unloading of the injured left hindlimb and compensatory loading of the contralateral limb.

## 3 Discussion

In this study, we developed a scalable tactile sensing platform and, as a first implementation, a tactile runway that measures how rodents dynamically distribute their load on the ground while walking freely. With 16,384 taxels at 2-mm pitch across a 512 *×* 128 mm surface and a nominal 1-kHz acquisition rate, the platform resolved locomotion at three scales simultaneously: movement along the runway, load distribution among the four paws and pressure dynamics within each footstep. Compared with optical footprint systems, which infer loading indirectly [13, 14, 17], and with force plates and pressure walkways [1, 21], the platform combines direct loading measurement with millimeter-scale resolution and kilohertz-rate acquisition, enabled by an adaptive readout that samples a fixed budget of taxels around active contacts in each frame [29] and a modular architecture in which spatial coverage and frame rate can be traded off in software. This allowed gait to be analyzed as a continuous process of weight support and transfer throughout stance. Because the hardware designs, software and recordings are openly available, the platform can be reproduced and adapted by other laboratories.

The sciatic nerve pinch model illustrates what direct contact measurement adds: measuring how much weight each paw carried, and not only how long it stayed on the ground, revealed deficits that step timing alone understated. Early after sciatic nerve pinch, the left-to-right hindpaw duty-factor ratio fell by 0.33 relative to baseline, whereas the corresponding time-integrated loading ratio fell by 0.59 (Fig. 5d): the rats continued to place the injured paw but bore much less weight on it. The loading data also showed how weight was redistributed during recovery. As the peak loading of the injured left hindpaw decreased, that of the contralateral hindpaw increased (Fig. 5e,f), and hindpaw imbalance shifted toward right-sided loading in all three SNI rats (Fig. 5h). Injured-paw contacts also became smaller and more elongated (Fig. 5c), indicating a change in how the paw was placed on the surface. By days 11–13, the duty-factor and loading ratios had moved back toward baseline. These observations are consistent with compensatory unloading described with pressure-mapping walkways [1] and altered paw contact after sciatic nerve injury [17, 32]. Because each contact was tracked from initial touch-down to lift-off (Fig. 4a; Supplementary Fig. S3), paw-specific dynamics could also be followed within stance. Paw-specific COP trajectories suggested delayed anterior load transfer in the injured hindpaw, and in SNI rats the net COP displacement at lift-off shifted by up to about 1.4 mm after surgery, mainly in the contralateral hindpaw (Supplementary Fig. S3). These features remain exploratory given the small sample size, but they illustrate a class of question that requires resolving load within stance: contacts can differ in when load is transferred, not only in how far the COP travels. Together, these results show that paw-resolved loading captured both the deficit in the injured limb and the compensation by the contralateral limb over the course of recovery.

This study has several limitations. Calibration was performed on board-level summed responses rather than on individual taxels, and held-out errors of 23.4–30.6 g (roughly 5–8% of body mass) indicate that absolute loading estimates remain approximate. These errors would be prohibitive for mice, whose body mass of roughly 25 g is comparable to the error itself, and the 2-mm taxel pitch would resolve little detail within a mouse paw. Adapting the platform to mice would therefore require finer-pitch sensor modules, which the modular design can accommodate, together with higher amplifier gain, which is set in software through the programmable digital potentiometer, and recalibration over a correspondingly lower load range. Combined with overhead video, automated paw assignment correctly labeled 93.3% of manually annotated contacts within curated walking windows but mislabeled 4.2% and missed 2.6%, so the biological analyses relied on manually corrected labels. Walking-window selection and camera-to-sensor registration also required manual curation, and automating these steps will be necessary for unsupervised analysis of longer sessions or larger cohorts. Finally, the cohort of three SNI and two sham rats was designed to demonstrate feasibility for longitudinal study, not to establish the specificity or reproducibility of each observed change, and walking speed may have contributed to some differences [33]. Extending the approach to turning, overlapping contacts and prolonged spontaneous behavior will require broader validation.

The next opportunity is to extend these measurements to other injuries and behaviors. Weight-bearing is an important characteristic to investigate in many rodent models, including spinal cord injury, osteoarthritis and peripheral nerve repair [4, 24, 30], and paw-resolved loading could test whether within-contact load transfer distinguishes injury types or tracks recovery earlier than timing-based measures. Because each board samples a fixed budget of taxels per frame (200 of 4,096 in this study), the trade-off between spatial coverage and frame rate, as well as sensitivity through programmable amplification, can be retuned in software for each application, provided each configuration is calibrated. Our sampling analysis indicates how acquisition rate should be set: for stances of 0.2–0.5 s, cumulative-loading error remained below 2% down to 50 Hz but rose to about 10% at 10 Hz (Fig. 4e), so the rate should be matched to contact duration and the measure of interest, with higher rates likely to matter most for briefer contacts. Jumping, for example, compresses take-off and landing into brief contacts with loads that can exceed body weight; a dedicated take-off and landing platform, recalibrated for that loading range, could quantify how animals push off and absorb impact before and after injury. Because each module has its own readout board, the sensing surface can also be extended to larger or differently shaped arenas; the same four modules can be tiled 2 *×* 2 into an open-field arena that records single and multiple freely moving rats (Supplementary Video 5), and extending the analysis pipeline to such recordings would enable studies of how social interactions influence locomotor coordination and paw loading.

Combining dense contact measurements with complementary mechanical and neural measurements is a further step. Shear sensing at comparable spatial resolution would allow normal and tangential loading to be studied together within a paw, while synchronized neural and muscle recordings could connect these mechanical events to sensorimotor control. Proprioceptive feedback contributes to locomotor coordination and recovery [34, 35], and somatosensory cortical activity is associated with locomotion in freely moving mice [36]. Resolving when and where each paw loads the ground would provide a mechanical reference for those signals and, with validated low-latency synchronization, could support perturbations triggered at specific phases of contact. More broadly, scalable tactile sensing could make physical interaction with the environment a routinely measurable component of freely moving behavior, complementing the increasingly sophisticated tools available for observing movement and neural activity.

## 4 Methods

### 4.1 Pressure-sensing platform and synchronized acquisition

The gait platform consisted of a high-density piezoresistive pressure-sensor array positioned beneath a protected walking surface, four independent readout boards, and two video cameras, one is placed overhead and the other one is placed on the side to assist manual paw labeling. Four 64 *×* 64 sensing modules were assembled into a 64 *×* 256-pixel runway array containing 16,384 tactile pixels at a 2-mm pitch. A waterproof protective cloth is laid flat on the sensors to prevent direct contact between the animal and the electronics while preserving pressure information from contact. Assembly of the runway is shown in Supplementary Video 1. Design files for the sensor module and readout board (KiCad sources, fabrication files and bill of materials), the firmware and a build guide are available online (see Code availability).

A custom Python program configured the four readout boards and recorded pressure and video data simultaneously. Each board (Teensy 4.1) was connected to the host computer through native USB serial. Before each trial, the host transmitted the maximum number of measurements retained per pressure frame, *M*, the intensity threshold, *T*, and the digital-potentiometer setting.

Rather than scanning all 4,096 taxels, each board measured a fixed budget of *M* taxels per frame (*M* = 200, approximately 4.9% of the module) and transmitted only those measurements. Taxels that had triggered neighbourhood sampling in the previous frame were measured first, so that ongoing contacts were tracked between frames. The remaining budget was spent on a background scan that visited taxels in coarse-to-fine order, beginning at the module centre and continuing through grids of successively halved pitch (32, 16, 8, 4 and 2 taxels), so that an incomplete scan still covered the module uniformly. The scan pattern was shifted by one column each frame, and by one row after each full sweep of columns, so that all taxels were measured across successive frames. Whenever a measured value exceeded *T*, the unmeasured taxels in its 3 *×* 3 neighbourhood were measured, and this was repeated recursively for any neighbour that also exceeded *T*, so that each contact was captured at full resolution. Neighbourhood measurements drew on the same per-frame budget, so frames containing contacts reached a coarser level of the background scan. Each taxel was measured at most once per frame.

Each board transmitted binary packets containing its identifier, frame number, and the indices and digitized values of the recorded taxels. Measurements were stored as a sparse indexed representation of each 64 *×* 64 pressure frame and were reconstructed into dense pressure maps.

Pressure and video recording threads were released using a common software start signal. Pressure packets and video frames were timestamped relative to the same host-clock reference, enabling synchronization. The cameras recorded at 1920 *×* 1080 pixels and a target rate of 30 frames per second.

### 4.2 Sciatic nerve pinch surgery

Six male Lewis rats (12–16 weeks old, 390–430 g) were included under protocol RA24M61. Four rats underwent left SNI and two underwent sham surgery; one SNI rat died after postoperative day 3 and was excluded (see Statistics and reproducibility). Anesthesia was induced and maintained with isoflurane (induction 3% isoflurane in oxygen at 0.5 L/min; maintenance 2% at 0.5 L/min). Depth of anesthesia was verified by absence of the toe-pinch withdrawal reflex. Rats were placed in lateral recumbency, and the operative hind limb (left) was shaved and aseptically prepared with betadine scrub. The lateral mid-thigh region, between the greater trochanter and the knee, was identified as the surgical field.

A 1–2 cm longitudinal skin incision was made over the lateral aspect of the thigh, centered over the biceps femoris. The fascia was incised, and the fibers of the biceps femoris muscle were bluntly separated to expose the sciatic nerve running posterior to the femur at mid-thigh level. Perineural connective tissue was carefully cleared to visualize the full circumference of the nerve without the nerve being stretched or dried.

An SNI was produced at this mid-thigh site using sterile thumb forceps applied perpendicular to the nerve trunk and held with consistent manual force for 20 s. The muscle was allowed to fall back into place, and the skin was closed using 4-0 interrupted sutures. At the end of the procedure, rats received buprenorphine (1.2 mg/kg, IM) for analgesia and were monitored during recovery from anesthesia until regaining sternal recumbency and spontaneous ambulation, then observed at least daily for signs of pain, infection, or wound dehiscence according to predefined endpoints. Sham rats (n = 2) underwent the same general surgical handling without nerve pinch injury.

### 4.3 Pressure-sensor characterization

The tactile array was characterized with an independent force plate while covered with the same waterproof layer used during animal experiments. Measurements were acquired from a 64 *×* 64 module using the same hardware configurations as for the animal experiment.

Dynamic response was assessed using four consecutive presses recorded simultaneously by the tactile array and force plate. The spatially integrated tactile response was interpolated to the force-plate timestamps and smoothed using a five-sample moving average. Active press intervals were defined as contiguous periods surrounding each tactile peak during which the signal exceeded baseline by either three baseline standard deviations or 3% of the peak response, whichever was greater.

The force-plate and tactile signals were independently min–max normalized, and agreement during the active intervals was quantified using

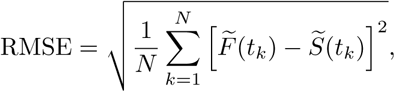

where *F̃* (*t_k_*) and *S̃*(*t_k_*) denote the normalized force-plate and tactile responses. This analysis assessed the agreement of normalized temporal responses rather than absolute calibration, which was evaluated separately in grams as described below. One representative press was divided into 20 temporal regions to visualize the evolution of the spatial pressure profile.

#### Board-specific calibration and held-out-press evaluation

The single-module examples above were complemented by separate calibration curves for all four boards (Supplementary Fig. S1d). Each curve used paired frame sums and reference loads from that board’s press recording. The first 3 s of each recording were excluded to avoid the start-up transient, corrupted readings (values outside the 12-bit ADC range) were discarded, and tactile frame sums were linearly interpolated to the force-plate timestamps. The force-plate baseline was estimated as the median of the lowest 20% of readings and subtracted from the reference signal.

Samples with reference loads of 0–450 g were sorted by load and divided into 20 approximately equal-count groups. Median taxel sums and reference loads defined calibration points, together with a zero-load baseline estimated from samples between *−*3 and 3 g. Monotonicity was enforced and duplicate taxel-sum points were removed, leaving 17–19 points per board. Piecewise cubic Hermite interpolation (PCHIP) [37] converted taxel sums to mass-equivalent loading. The exported curves ended at approximately 410–421 g, depending on the board. Application to runway contacts is described in Section 4.8.

Performance was evaluated by leaving out one entire selected press at a time and refitting the calibration on the remaining presses (Supplementary Fig. S1e). Presses were separated at minima between detected force peaks, with each boundary sample assigned to only one press. Reference tare, sensor baseline and calibration points were estimated from training presses only; the training tare was also applied to the held-out reference. Predictions used PCHIP. Evaluation included held-out reference samples between 0 and 450 g. RMSE pooled the scored time samples across folds, including near-zero loads. This evaluation used the existing selection of presses within each recording and did not test transfer to independent sessions or different contact geometries.

### 4.4 Force-plate calibration and acquisition

The reference force platform consisted of three 5 kg full-bridge load cells wired in parallel and connected to a Teensy microcontroller through a 24-bit NAU7802 amplifier. The load-cell bridge was excited at 3.3 V, with an amplifier gain of 128 and a conversion rate of 320 samples per second.

Calibration measurements were collected with the unloaded platform and with known masses spanning the operating range. Each calibration point was obtained by averaging 512 consecutive ADC readings. A least-squares linear model,

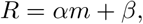

was fitted between the averaged raw output *R* and applied mass *m*, where *α* is the calibration scale and *β* is the zero-load offset. During characterization experiments, each reported force-plate measurement was averaged over 32 ADC conversions, producing an effective output rate of approximately 10 Hz.

### 4.5 Pressure-map reconstruction and video preprocessing

Pressure packets from the four boards and frames from the overhead cameras were synchronized using their shared timeline. At each time point, the most recent valid packet from each board was used to reconstruct a local 64 *×* 64 pressure image. Taxel indices were converted to row and column coordinates, board-specific orientation corrections were applied, and the four images were concatenated into a 64 *×* 256 runway pressure map.

Candidate paw contacts were detected from the reconstructed pressure maps. Each pressure frame was thresholded to generate a binary contact mask, which was morphologically opened and hole-filled before connected-component analysis. Nearby pressure components were merged when their spatial separation and combined area were consistent with a single paw contact. Small, dispersed, or low-intensity components were excluded using area, density, and mean-pressure criteria.

Overhead video frames were processed independently to extract the animal body mask. For each recording, a background frame was generated by taking the pixel-wise median of sampled video frames. Foreground pixels were identified by background subtraction, and the resulting binary mask was refined using morphological closing, opening, and hole filling. Small components were removed, and the largest, valid remaining component was retained as the animal body mask.

### 4.6 Tail-base localization, camera-to-tactile registration and body orientation

Before registered fusion with the pressure data, a lightweight convolutional encoder–decoder localized the tail base in the overhead video. This landmark resolved the anterior–posterior ambiguity of the body major axis. The task-specific model estimated a single landmark using the body mask to constrain the relevant image region. The network received a four-channel input consisting of the RGB overhead image and binary body mask and produced a single-channel heatmap at the original image resolution, as in heatmap-based pose estimation [9, 10].

Manual tail-base annotations were converted into Gaussian target heatmaps. For an annotated tail-base coordinate (*x_i_, y_i_*), the target heatmap was defined as

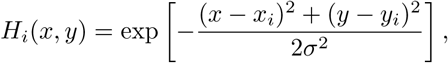

where *σ* is a fixed hyperparameter that controls the width of the Gaussian target heatmap and therefore the spatial tolerance around the manual annotation. The network was trained using pixel-wise mean squared error between the predicted heatmap, *H*^^^*_i_*(*x, y*), and the corresponding target heatmap:

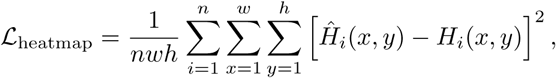

where *n* is the number of training images and *w* and *h* are the heatmap width and height, respectively. During inference, the predicted tail-base coordinate was obtained from the maximum of the output heatmap,

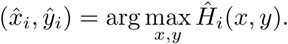

Detector performance was assessed by 5-fold cross-validation across 73 recordings, holding out whole recordings and selecting the training epoch on separate recordings, repeated 5 times (results in Section 2.6).

Camera coordinates were aligned with the tactile array using a projective transformation. For each recording, the four corners of the platform were manually identified in an overhead video frame. A reference template, defining the position of the sensor sheet relative to the platform, was used to obtain the corresponding sensor-array corners, which were matched to the corners of the tactile grid to calculate a perspective transformation mapping video coordinates to sensor coordinates.

The body mask and predicted tail-base location were projected into tactile coordinates. The projected body centroid and tail-base location established body orientation, allowing the camera-guided detector to accumulate FL, FR, HL and HR votes for each contact. These votes supplemented the pressure-based assignment, whose primary reference was the walking direction estimated from smoothed COP motion.

### 4.7 Pressure-based contact tracking and camera-assisted paw assignment

Candidate walking windows were automatically extracted based on whole-animal COP speed followed by manual curation. Pressure-based contact tracking was combined with saved camera-derived paw votes. This automatic pipeline was evaluated separately from the manually reviewed labels used for longitudinal analyses.

#### Paw-assignment pseudocode

**Input:** Pressure frames, selected walking windows and saved camera-derived paw votes.

**Output:** Stance events labeled FL, FR, HL or HR.

1. Calculate whole-animal COP and smooth its trajectory.
2. For each selected walking window:

(a) Estimate the window’s overall travel direction from COP displacement.
(b) Detect contact regions in each pressure frame and track them across successive frames.
(c) Join nearby fragments. Split tracks at significant loading minima to separate successive contacts from different paws that happen to land at nearby locations, and discard contacts below duration or local intensity thresholds.
(d) For each contact, at its stance midpoint:

(i) Estimate local heading and contact offsets from smoothed COP.
(ii) Calculate assignment scores for each of the four paws using the contact’s position relative to the smoothed COP and walking direction.
(iii) Retrieve paw votes from the nearest overlapping camera-guided detector event, if available.
(iv) Combine position scores with weighted camera-vote scores.
(e) Group nearby overlapping fragments and sum their scores.
(f) Combine left/right evidence to choose forepaw or hindpaw identity.
(g) Within each forepaw or hindpaw sequence, decode left/right labels in onset order using Viterbi decoding [38]; penalize consecutive overlapping contacts assigned to the same side.
3. Retain contacts with stance midpoints in the selected walking windows, remove duplicate fragments, and merge consecutive nearby pieces assigned to the same paw.

#### Tracking parameters

Taxels with values above 200 raw ADC counts were identified as active. A 3 *×* 3 binary dilation connected nearby active regions before individual contact regions were identified. Linking allowed 4-taxel (8-mm) displacement and three missing frames (about 3 ms); fragment joining allowed 4 taxels (8 mm) spatial displacement and 50 ms temporal separation. Continuous tracks containing multiple loading peaks were recursively split at minima below 50% of the smaller peak on either side, helping separate successive contacts from different paws that would otherwise be grouped as a single event. Contacts were retained if they lasted at least 0.10 s and their peak summed taxel response reached 4 *×* 10^4^ summed ADC counts (approximately 18–29 g mass-equivalent, depending on the board). Tracking extended 0.5 s beyond walking-window boundaries.

#### Reference frame and scoring

COP used the same threshold, with linear interpolation of missing samples and boundary extrapolation. Longitudinal and lateral references used 0.3-s and 0.6-s moving averages. Heading followed the 0.3-s-smoothed COP displacement over 1.5 s centered on the stance midpoint, falling back to the walking window’s overall travel direction along the runway axis when displacement was *≤* 3 taxels (6 mm) or opposed that direction.

At the midpoint of each stance interval, the contact’s position relative to the smoothed COP was measured along and perpendicular to the estimated walking direction. These offsets, denoted by *a_j_* and *ℓ_j_*, were expressed in taxels, with positive values indicating forward and rightward displacement, respectively. Assignment scores were calculated using the logistic sigmoid function, *σ*(*z*) = (1 + *e^−z^*)*^−^*^1^:

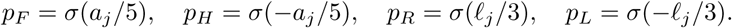

The 5- and 3-taxel (10- and 6-mm) scale parameters controlled the steepness of the longitudinal and lateral assignment transitions, respectively. A smaller lateral scale produced a sharper transition between left and right assignments, reflecting the closer spacing between contralateral paws than between forepaws and hindpaws.

For contact *j*, camera-derived votes were obtained from the nearest temporally overlapping detector event within 8 taxels (16 mm). Let *v_j,k_* denote the number of votes for paw identity *k ∈* {FL, FR, HL, HR}. The camera support for each paw was

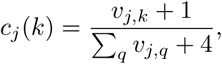

where the sum runs over all four paw identities. Adding one vote per identity prevented zero probabilities; when no votes were available, *c_j_*(*k*) = 1*/*4 for every paw.

Position and camera information were combined as

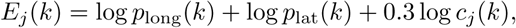

where *p*_long_(*k*) is the corresponding forepaw or hindpaw score, and *p*_lat_(*k*) is the corresponding left or right score. The coefficient 0.3 reduced the influence of camera-derived evidence.

#### Temporal constraints

Overlapping fragments less than 5 taxels (10 mm) apart shared summed scores, with the longest retained. Viterbi decoding penalized consecutive same-side assignments within each forepaw or hindpaw sequence by log(0.01) when overlap exceeded 20 ms, without enforcing alternation. Final same-paw merging allowed 6-taxel (12-mm) separation and gaps up to 60 ms.

#### Evaluation

Manual labels comprised 2,469 contacts from 81 recordings within selected walking windows. Annotators labelled contacts from the pressure data and the primary overhead video, and consulted the second camera when paw identity was ambiguous. Each reference contact was matched independently to the nearest automatic contact within 15 taxels (30 mm) whose interval contained its stance midpoint with a 50-ms tolerance; unmatched references were counted as missed. Matches were not constrained to be one-to-one. Agreement was 93.3% across all 2,469 reference contacts and 95.7% among the 2,406 matched contacts, excluding 63 missed contacts. Confusion-matrix rows include misses and are normalized by manually labeled paw counts (Supplementary Fig. S2d). This analysis scores reference contacts, not the precision of all automatic detections; repeated contacts and recordings are not independent animal replicates.

### 4.8 Pressure-derived gait metrics

For contact *j*, 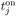 and 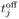 denote contact onset and lift-off. Stance duration, stride duration and duty factor were

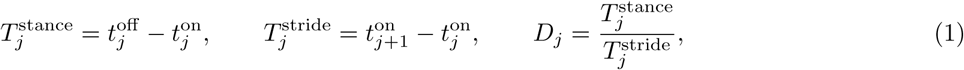

where *j* + 1 denotes the next contact of the same paw within the recording. Duty factor is the fraction of a stride spent in stance.

The summed tactile response for each contact was

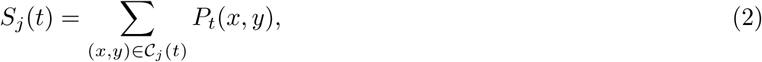

where *C_j_*(*t*) is the set of taxels assigned to contact *j* at time *t*, and *P_t_*(*x, y*) is the raw response of the taxel at position (*x, y*). Raw peak loading and time-integrated response were defined as

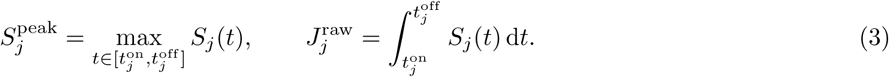

Contact area was calculated from the binary contact mask:

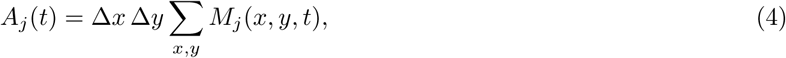

where *M_j_*(*x, y, t*) equals one for included taxels and zero otherwise. The taxel dimensions were Δ*x* = Δ*y* = 2 mm, giving an area of 4 mm^2^ per taxel. For morphology analysis, the maximum area during stance, 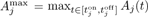, was calculated using the mask defined in the pressure morphology subsection.

Each runway contact was assigned to a sensor board using the horizontal centroid stored with the extracted contact event. The corresponding board-specific calibration curve converted the summed tactile response to massequivalent loading:

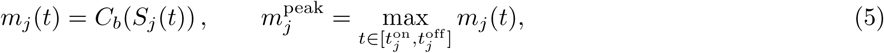

where *C_b_* is the calibration curve for board *b*, and *m_j_*(*t*) is expressed in grams. Raw measurements were retained alongside the calibrated values and a flag indicating whether the input fell within the calibration range.

Time-integrated loading was calculated by calibrating each sample before integration. For *n_j_*samples acquired during contact *j*, the composite trapezoidal rule gave

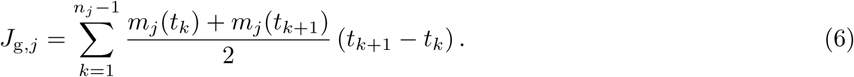

Thus, the mean loading between each pair of consecutive samples was multiplied by their time interval and summed over stance. The resulting integral has units of g s. Multiplication of mass-equivalent loading by 9.80665*×*10*^−^*^3^ N g*^−^*^1^ converts it to normal force in newtons; the corresponding time integral has units of N s.

For longitudinal comparisons (Fig. 5e,f), calibrated peak loading was normalized by body mass:

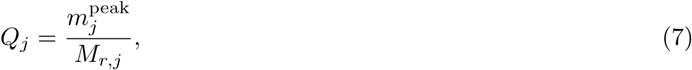

where *M_r,j_* is the body mass in grams of rat *r* for the recording containing contact *j*. For each rat, paw and epoch, the median *Q_j_* was divided by the corresponding presurgical median and multiplied by 100. Rat–paw–epoch summaries required at least five eligible contacts, and group curves were means of these individual-rat percentages.

Within each walking window *w*, left-to-right hindpaw ratios were calculated from the median duty factor and time-integrated loading:

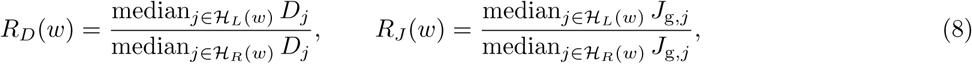

where *H_L_*(*w*) and *H_R_*(*w*) contain the left and right hindpaw contacts within that window, respectively; each window required at least two strides per hindpaw. For Fig. 5d, each rat’s epoch value was the mean of its window ratios in that epoch and required at least two windows. Group traces are the mean of these per-rat values, so each rat contributes equally regardless of how many walking windows it has.

Loading imbalance was evaluated separately during hindpaw-only support. These windows lasted at least 0.2 s, with both hindpaws in stance and neither forepaw in contact, according to the manually corrected labels. No locomotion criterion was applied. Windows were excluded if either hindpaw center was within 20 mm of the array edge.

In each support window, the left and right hindpaws were the labelled HL and HR contacts in stance, each represented by its tracked contact region; taxels falling in both regions were assigned to the nearer hindpaw centre. The imbalance index was

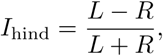

where *L* and *R* are the raw sensor responses summed over the HL and HR contact regions, respectively, and over all retained frames in the window, so that other contacts, such as the tail, did not contribute. Positive values indicate greater left-sided loading, and negative values indicate greater right-sided loading. Frames in which the hindpaws formed a single connected pressure cluster were omitted; windows were excluded if more than half of their frames were omitted. Session values were unweighted means of the window indices, and epoch values were means of the session values (Fig. 5g,h).

The center of pressure (COP) was calculated as the response-weighted centroid of a selected set of taxels *C*(*t*):

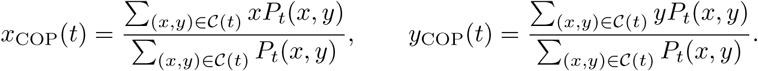

COP was defined only when the summed response was nonzero. For paw-level trajectories, *C*(*t*) contained the taxels belonging to an individual contact. For whole-animal trajectories, it contained the recorded surface contacts across the array. Whole-animal COP was compared with the video-derived body centroid (Fig. 3). Longitudinal COP position and COP speed described the position and motion of the pressure-weighted centroid, reflecting the distribution of surface loading rather than body center of mass.

### 4.9 Sampling-rate analysis

Manually reviewed stance events from all four paws were grouped by contact duration for the sampling-rate analysis. Paw-specific loading traces were reconstructed at native timestamps and downsampled to 500, 100, 50, and 10 Hz by retaining the native sample closest to each target timestamp. Downsampled traces and cumulative loading were compared with the native recording. Illustrative contacts from all four paws are shown in Fig. 4f.

### 4.10 Pressure morphology analysis

Footprint morphology was measured on manually labelled contacts within walking windows; contacts were included if they lasted 0.08–1.5 s, spanned at least five frames and were not fused with another labelled contact. Pressure frames were filtered with a three-frame per-taxel temporal median to remove single-frame readout artefacts, and the paw region was tracked from the labelled contact centroid in every stance frame. Fragments whose centres lay more than 20 mm from that centroid (29 mm for HL, whose injured paw often landed as separate islands), or closer to another labelled contact in stance, were removed. For each contact, the frame with the largest paw area was selected, and all paw taxels at or above 10% of that frame’s maximum, including separated pads, were used to measure contact area and the major and minor axes of the ellipse with the same second moments (length and width). Rat–epoch–paw cells with fewer than five contacts were excluded.

### 4.11 Statistics and reproducibility

This feasibility study used six rats; no statistical method was used to predetermine sample size. One SNI rat died after postoperative day 3 and was excluded, leaving three SNI and two sham rats. To avoid weighting sessions by their number of observations, each measure was averaged per recording session (log-transformed for the length/width ratio and peak loading). Within each group, session means were analyzed with linear mixed-effects models and a random intercept for rat, which allows each rat to have its own baseline level; each postoperative epoch was compared with presurgical baseline using two-sided tests with Satterthwaite degrees of freedom [39], without multiple-comparison adjustment. Inclusion thresholds and sample sizes are given in the figure legends and Methods.

## 5 Data availability

All 85 runway recordings analysed in this study are available on Zenodo at https://doi.org/10.5281/zenodo.23286364 (CC BY 4.0). Each recording includes the raw pressure packets with camera frame timestamps (MATLAB format) and the overhead video (MP4). The deposit also includes the original annotations (camera-to-sensor homographies, manual paw labels for 81 recordings, walking windows, reviewed contacts and body mass) and a demonstration package with one complete recording. Processed outputs are not deposited; they can be regenerated with the code described in Code availability. The Supplementary Videos are also available at https://github.com/tactilearena/tactilearena.github.io/releases/tag/supplementary-videos-v1. Other data are available from the corresponding authors upon reasonable request.

## 6 Code availability

The readout firmware, synchronized pressure and video acquisition software, and the analysis pipeline (pressure reconstruction, contact detection, paw labelling and gait-metric extraction, with a pretrained tail-base localization model, calibration data and a demonstration workflow with reference outputs) are available at https://github.com/tactilearena/tactilearena.github.io under the MIT License. Scripts used to compute the gait metrics and statistical analyses reported here are available from the corresponding authors upon request. The same repository contains the hardware design files for the sensor module and readout board (KiCad sources, fabrication files and bill of materials) under the CERN Open Hardware Licence v2 (permissive). Documentation, a build guide and the Supplementary Videos are also available on the project website, https://tactilearena.github.io.

## 7 Acknowledgements

The authors thank Rachana Suresh, MBBS, MPH, for her support of this research through the Johns Hopkins Merkin Peripheral Neuropathy and Nerve Regeneration Center Microgrant program. All animal procedures were conducted under a protocol approved by the Johns Hopkins University Animal Care and Use Committee (ACUC; protocol RA24M61, approved March 11, 2024). Artificial intelligence (AI) tools were used to assist with manuscript editing, including improvements to language, clarity, and organization. The authors reviewed and verified the final manuscript and take full responsibility for its content.

## 8 Funding Statement

This work is supported by the Johns Hopkins Merkin Peripheral Neuropathy and Nerve Regeneration Center, 2025 Microgrant. This work is supported by the Rita Levi Montalcini Fellowship offered by the Dompé Foundation in Year 2025.

## 9 Author contributions

W.L. programmed and implemented the experimental use of the pressure-imaging platform, designed and implemented the video and data-analysis pipeline, contributed to the experimental design, performed the animal experiments, analyzed and interpreted the experimental data, and wrote the original manuscript draft. A.S. conceived the original project, designed, developed, and validated the large-area pressure-imaging platform, and demonstrated its proof of concept; A.S. also contributed to the experimental design, interpretation of results, and manuscript preparation. P.P. performed surgical procedures and contributed to manuscript preparation. A.A.O. and K.Q. conceived the original project, contributed to the experimental design and manuscript preparation. N.T. supervised the project. All authors reviewed and approved the final manuscript.

## Competing interests

The authors declare no competing interests.

## Supplementary figures

**Fig. S1.**
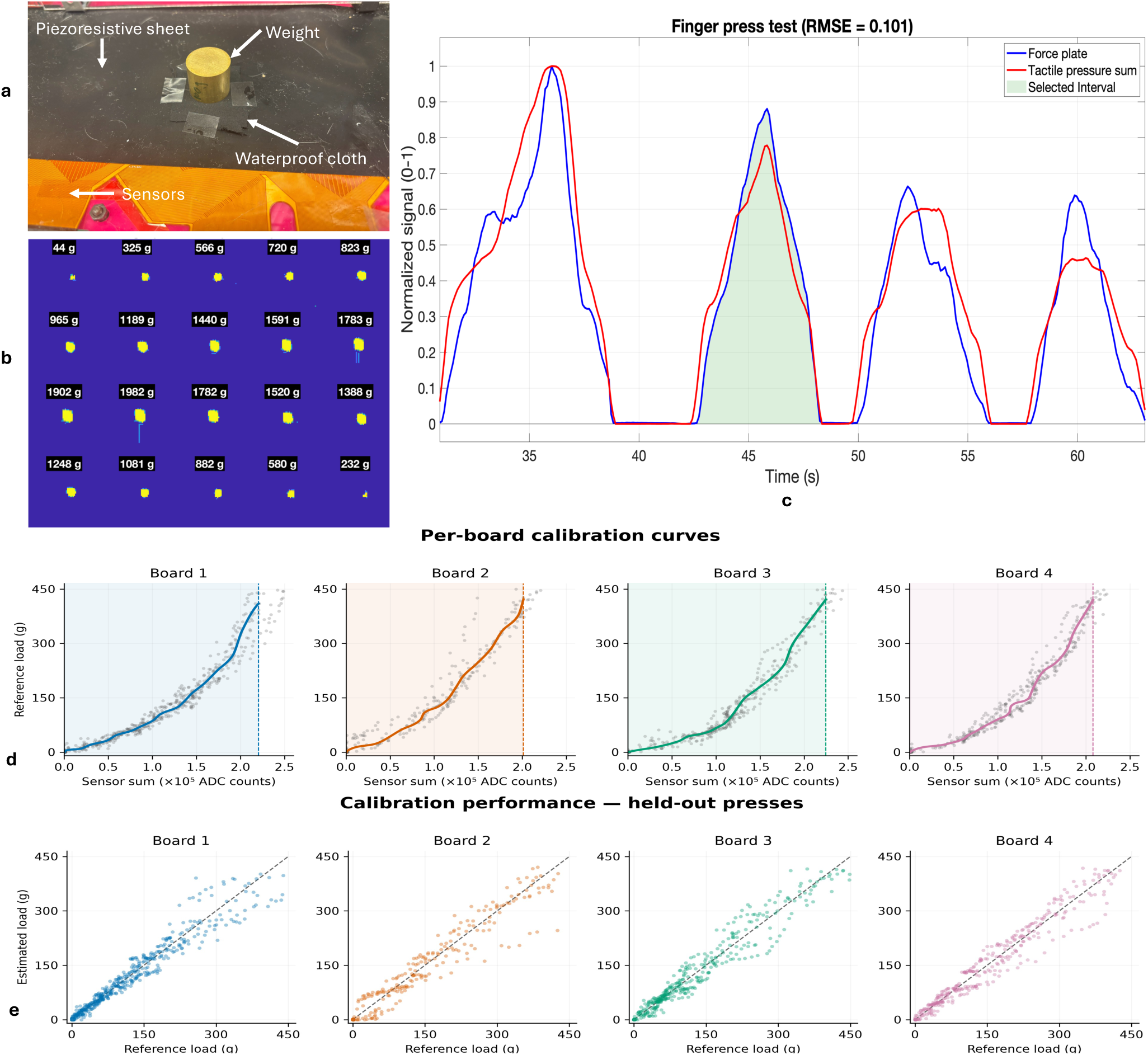
Sensor characterization and board-specific calibration. **a,** Characterization setup with a known weight placed on the waterproof layer covering the sensor. **b,** Twenty pressure-map snapshots spanning a representative press; labels give mass-equivalent readings from the reference force plate, and the color scale shows recorded taxels values. **c,** Independently normalized force-plate and summed tactile responses during four consecutive presses. The shaded interval corresponds to the snapshots in **b**; RMSE = 0.101 over the active press intervals. **d,** Calibration curves for boards 1–4. Gray points show selected paired taxels sums and reference loads; colored curves show PCHIP interpolation of the board-specific calibration points. Shading and vertical dashed lines identify the usable taxelsum domains. Loads are expressed in mass-equivalent grams. **e,** Estimated versus reference load for held-out indenter presses, evaluated by leave-one-press-out cross-validation within each board’s recording. Points represent paired time samples, and dashed lines indicate equality. RMSEs across scored samples were 23.4, 30.6, 25.9 and 27.2 g for boards 1–4, respectively.

**Fig. S2.**
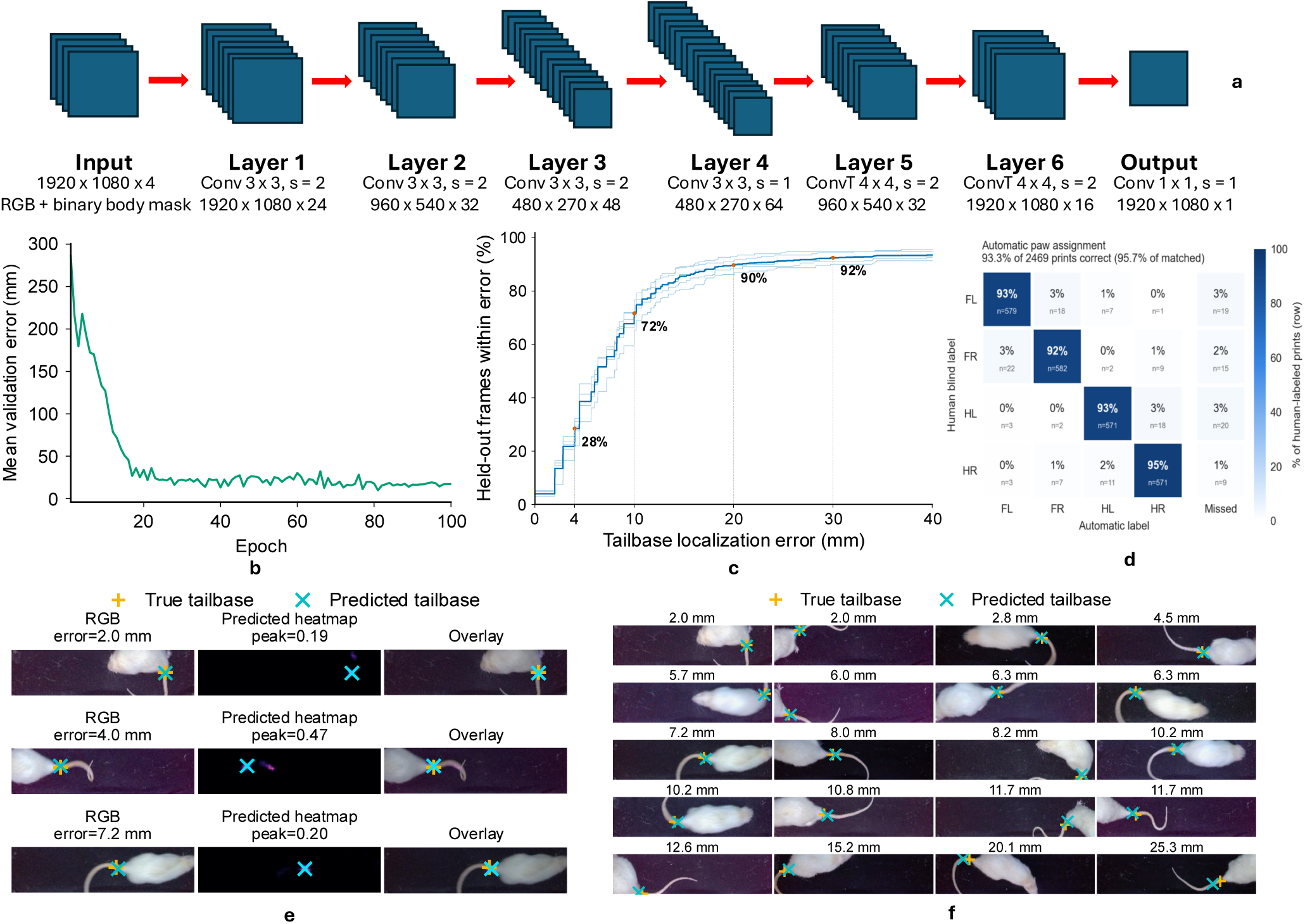
**a,** Encoder–decoder architecture using an RGB frame and binary body mask as input and producing a single-channel tail-base heatmap. **b,** Tail-base detector training curve: mean validation error across epochs. The lowest-error checkpoint (epoch 81; 9.8 mm) was kept; **c** gives an independent estimate. **c,** Fraction of held-out frames within increasing tail-base localization-error thresholds, in mm at the walking surface, from 5-fold cross-validation repeated 5 times; light lines show individual repeats and the dark line their average (see Section 2.6). **d,** Camera-assisted pressure-based paw assignment compared with manual labels for 2,469 contacts from 81 recordings within manually curated walking windows. Rows indicate manual labels and columns indicate automatic labels or unmatched reference contacts (Missed). Cells show counts and row percentages. The pipeline correctly labeled 2,303 contacts: 93.3% of all reference contacts and 95.7% of the 2,406 matched contacts. This panel evaluates the complete tracking and assignment pipeline, not the tail-base localizer alone. **e,** Example predicted heatmaps and overlays for frames with different localization errors. **f,** Representative examples comparing corrected and predicted tail-base locations across body postures. The number above each frame gives its tail-base localization error in mm; frames are ordered by increasing error.

**Fig. S3.**
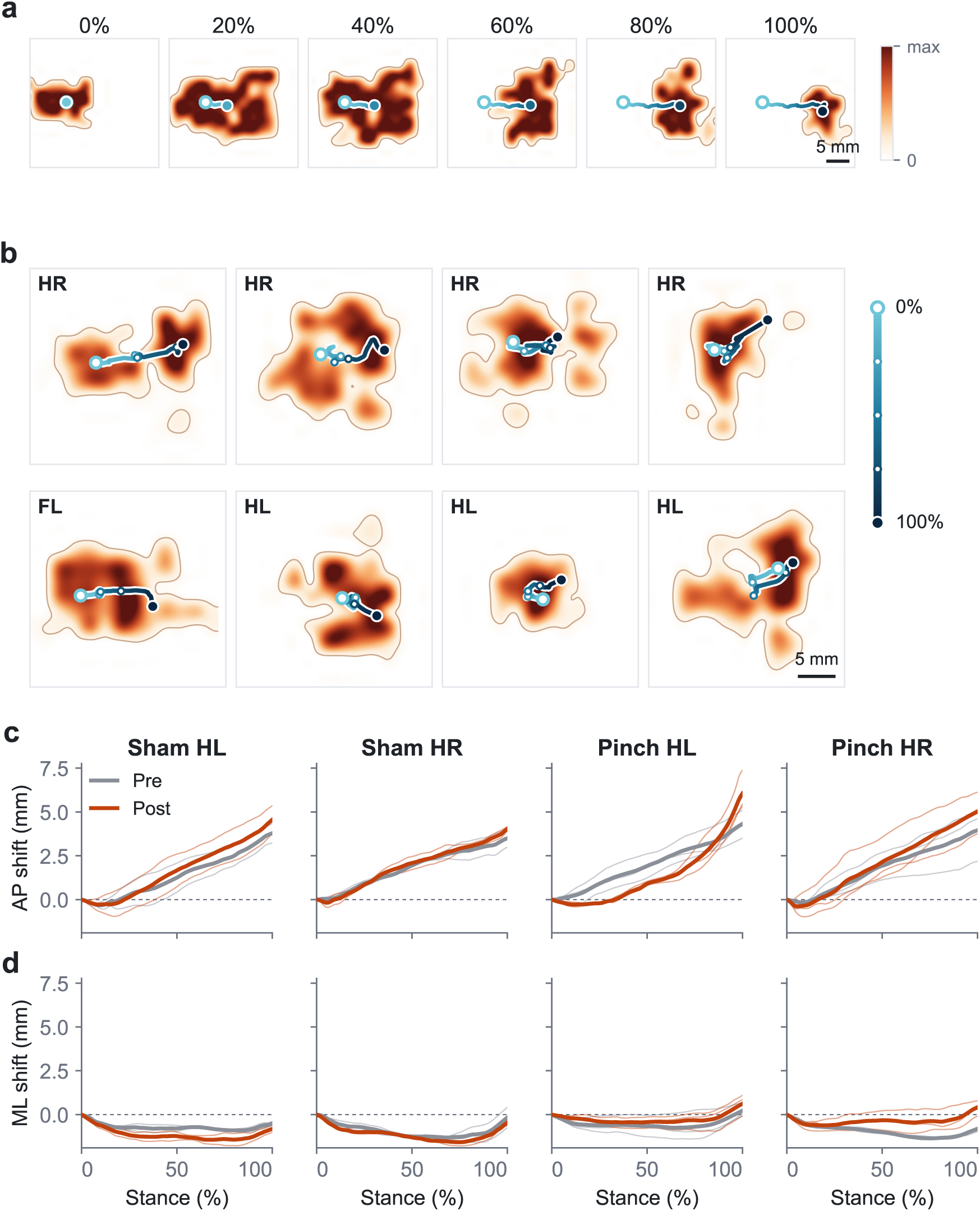
Within-paw COP trajectories. **a**, Pressure frames from one right hindpaw contact at 0–100% of stance, with the paw-level COP trajectory up to each frame. **b**, Representative contacts: pressure–time integral with the COP trajectory from initial contact (0%, open circle) to liftoff (100%, filled circle); small markers indicate 25, 50 and 75% of stance. Paws are mirrored so that the direction of travel is to the right. Scale bars, 5 mm. **c,d**, Hindpaw COP displacement from initial contact over stance in the anterior–posterior (**c**; positive toward the direction of travel) and medio-lateral (**d**; positive lateral) directions, before (pre) and after (post, days 1–13 pooled) surgery in sham (*n* = 2) and pinch (*n* = 3) rats, from 910 human-labeled walking prints. Thin lines, per-rat means; bold lines, mean of rats. Within each group, net displacement at liftoff was compared between presurgical and pooled postoperative recording-session means (linear mixed-effects model with a random intercept for rat; Satterthwaite degrees of freedom). In pinch rats, it shifted laterally by 0.7 mm in HL (*p* = 0.032) and anteriorly by 1.4 mm (*p* = 0.026) and laterally by 0.9 mm (*p* = 0.019) in HR, with no significant anterior–posterior change in HL (*p* = 0.24); sham rats showed no significant change (all *p ≥* 0.22). The within-stance COP shift in AP direction also indicates a timing variation that occurs after surgery: slower initial contact and sharper liftoff. Pressure maps are displayed with bicubic interpolation; COP was computed from the uninterpolated taxel data and smoothed with a 40 ms moving average. These data illustrate load transfer within individual contacts. Whole-animal trajectories are shown in Fig. 3.

## Supplementary Videos

**Supplementary Video 1 | Assembly of the tactile runway.** Each 64 *×* 64 sensor module is connected to its readout board through its two flexible tails; the four modules are aligned edge to edge, covered with a piezoresistive film, and connected to the host computer. On-screen captions describe each step.

**Supplementary Video 2 | Synchronized overhead video and paw-resolved plantar pressure during walking before sciatic nerve pinch.** Overhead video of a rat walking on the runway before surgery (rat 6, 11 February 2026), with each paw contact labeled by the analysis pipeline (FL, FR, HL, HR). The lower panels show the plantar pressure distribution of each paw during stance as surfaces of raw taxel values. Playback at 0.25*×* real time.

**Supplementary Video 3 | Synchronized overhead video and paw-resolved plantar pressure during walking after sciatic nerve pinch.** As Supplementary Video 2, nine days after left sciatic nerve pinch (rat 4, 19 March 2026). Playback at 0.25*×* real time.

**Supplementary Video 4 | Adaptive tactile sampling during locomotion.** First, one acquisition frame of a single 64 *×* 64 module, shown read by read: the readout firmware’s sampling loop applied to a simulated paw contact, which re-reads the previous frame’s active taxels, performs a coarse-to-fine background scan and recursively reads the 3 *×* 3 neighbourhood of every reading above threshold until *M* = 200 taxels have been read. Then, recorded packets from all four modules during walking (rat 4, 24 February 2026), displayed frame by frame and at approximately 1/8 of real time. Black, not read in that frame; grey, read and below threshold (*T* = 700); colour, read and above threshold.

**Supplementary Video 5 | Reconfiguration of the sensor modules into an open-field arena.** Four 64 *×* 64 modules tiled 2 *×* 2 form a 256 *×* 256 mm open-field arena of 128 *×* 128 taxels. First, a single rat walking, turning and rearing (rat 4, 24 February 2026; *M* = 200, *T* = 700), shown with the tactile map, an overhead camera and a side camera at 0.75*×* real time. Then, three rats exploring the arena simultaneously, shown with the tactile map and the overhead camera.

## Notes

https://tactilearena.github.io/

https://github.com/tactilearena/tactilearena.github.io

https://doi.org/10.5281/zenodo.23286364

### Competing Interest Statement

The authors have declared no competing interest.

### Summary of Updates

This version clarifies the methods, reports additional quantitative detail and improves the figures. No conclusions have changed. Physical units and scale are reported more explicitly throughout. Tracking tolerances are given in millimetres, detection thresholds in ADC counts with approximate mass equivalents, and the force-plate RMSE is placed in context against absolute calibration errors in grams. The Discussion now outlines the modifications needed to adapt the platform to mice: finer-pitch sensor modules, higher amplifier gain and calibration over a lower load range. The tail-base localizer is now evaluated by repeated five-fold cross-validation across 73 recordings. It achieved a median error of 6.6 mm, with 72% and 90% of frames within 10 and 20 mm, and these results are reported in Section 2.6. Section 2.4 notes that postsurgical walking windows were about four times longer than presurgical ones, consistent with reduced stride length. Figure captions now state the normalization of the pressure maps, the orientation of the filmstrips, axis ranges that differ between panels, and localization error in millimetres. Figure 5 has been rebuilt at final print size with legible labels. All figures were re-exported as vector graphics, and minor typographical errors have been corrected.

